# Revisiting the Association between Human Leukocyte Antigen and End-Stage Renal Disease

**DOI:** 10.1101/2020.03.18.996330

**Authors:** Naila Noureen, Farhad Ali Shah, Jan Lisec, Hina Usman, Mohammad Khalid, Rimsha Munir, Nousheen Zaidi

## Abstract

Multiple works have studied possible associations between human leukocyte antigen (HLA) alleles and end stage renal disease (ESRD) showing, however, contradictory and inconsistent results. Here, we revisit the association between ESRD and HLA antigens, comparing HLA polymorphism (at HLA-A, -B, -C, -DRB1, -DQB1 and DQA1 loci) in ESRD patients (n=497) and controls (n=672). Our data identified several HLA alleles that displayed a significant positive or negative association with ESRD. We also determined whether heterozygosity or homozygosity of the ESRD-associated HLA alleles at different loci could modify the prevalence of the disease. Few HLA allele combinations displayed significant associations with ESRD among which HLA-A*3–HLA-A*26 combination showed the highest strength of association (OR= 4.488, P≤ 0.05) with ESRD. Interestingly, the age of ESRD onset was not affected by HLA allele combinations at different loci. We also performed an extensive literature analysis to determine whether the association of HLA to ESRD can be similar across different ethnic groups. Our analysis showed at least for certain alleles, HLA-A*11, HLA-DRB1*11, and HLA-DRB1*4, a significant association of HLA to ESRD in different ethnic groups. The findings of our study will help in determining possible protective or susceptible roles of various HLA alleles in ESRD.

## Introduction

A wide array of research works have indicated associations between human leukocyte antigen (HLA) status and various kidney diseases [1]. Few of these disorders are immune-mediated; while, in others the pathogenesis is unclear or the relevance of HLA is not completely understood.

In particular, end-stage renal disease (ESRD) –the last stage of chronic renal failure– has become a global health problem [2] and was hence investigated for HLA association repeatedly [3-5]. Most of these studies have examined waitlisted candidates for renal transplantation. Nonetheless, there are several contradictions in these previous works and no consistent HLA associations with ESRD itself have been identified [1]. It has been suggested that few HLA alleles may affect the severity of kidney disease or risk of progression [1]. For instance, specific HLA alleles could promote a more generalized pro-fibrogenic T cell phenotype that may contribute to disease progression or onset [1].

The presented study aims to revisit the association between HLA-polymorphism and ESRD. Most of the previous works have studied this association in homogeneous ethnic groups and they have interpreted their data accordingly. For this study, we compare the HLA polymorphism (at HLA-A, -B, -C, -DRB1, -DQB1 and DQA1 loci) in ESRD patients and controls from Punjab –one of the better-developed regions in Pakistan. To the best of our knowledge, none of the previous works have studied the association between HLA alleles and ESRD in the Pakistani population. Also, only a few previous works have studied HLA allele frequencies in the normal Pakistani population [6-9] (**Supplementary Table S1**). These studies also have limitations with respect to the population size [6, 7] or the limited number of HLA loci studied. Here, we try to overcome these limitations by studying the broader-array of HLA genes in a larger population sample. Previous works have shown that for certain pathological conditions heterozygosity or homozygosity of the disease-associated HLA alleles could modify the manifestation of that condition [10]. Hence, here we aimed to determine whether heterozygosity or homozygosity of the ESRD-associated HLA alleles at different loci could modify the prevalence of the disease. Moreover, we also compared our data with major previous studies to identify any consistent patterns of HLA allele associations with ESRD. The findings of this study will help in determining possible protective or susceptible roles of various HLA alleles in ESRD.

## Materials and Methods

### Study-population, ethics and sample collection

For the present study, we examined the ESRD patients (n=497) that were waitlisted (from 2017-2019) for renal transplantation at various transplant centers across Punjab, Pakistan. In addition, healthy subjects (n=672) from the same region were also included in this study as a control population. All the study subjects were above the age of 15. We excluded subjects with the following conditions: severe viral/bacterial infection, on anticoagulation therapy, suffering from bleeding disorder (e.g. hemophilia, low platelets, etc.) and aplastic anemia. The study protocol for human subjects was approved by the Research Ethics and Biosafety Committee-MMG, University of the Punjab. Informed consent was obtained from each study-subject before sample collection. To obtain the basic personal information and medical history each participant was interviewed and completed a structured questionnaire. The medical history file of each patient was also thoroughly examined. Intravenous blood was collected from all the subjects according to the guidelines of the National Committee for Clinical Laboratory Standards (document H18-A4) [11] in vials containing EDTA-anticoagulant agent.

### DNA extraction and HLA genotyping

Genomic DNA was extracted from whole blood using DNAZOL BD reagent according to the manufacturer’s instructions (Thermofisher Scientific). DNA concentration and purity were determined by Nanodrop spectrophotometer (ThermoScientific, USA). The DNA samples were stored at −20 °C until use. The samples were typed for HLA-A, -B, -DRB1, -DQB1,-DQA1 loci by sequence-specific oligonucleotide PCR (SSO-PCR) using LIFECODES HLA SSO typing kits. For HLA-C typing LIFECODES HLA-C eRES SSO typing kits were used. The product signals were detected by Luminex-200 and XY platform and were analyzed by MatchIT DNA software (Immucor GTI diagnostics inc. USA) according to the instructions mentioned in the software manual.

### Statistical analysis

HLA-A, -B, -C, -DRB1,-DQB1, and DQA1 allele frequencies (AF) were determined for each allele in patients with ESRD and controls using the following formula: AF (%)=(n/2N)×100, where n indicated the sum of a particular allele and N indicated the total number of individuals. The differences in allele percentages between patients with ESRD and controls were analyzed by cross-tabulation using the Fisher test. In the statistical process, P-value was calculated according to the expected value. The strength of disease association to a particular allele was expressed by odds ratio (OR) at 95% confidence intervals (95% CI). Statistical significance was accepted when P < 0.05. Alleles with OR >1.00 were considered to be positively associated with ESRD. Alleles with OR < 1.00 were considered to be negatively associated with ESRD. Analysis of allele combination was performed using the R software environment 3.4.2 (http://cran.r-project.org/).

## Results

### HLA-A, -B, -C, -DRB1, -DQB1 and DQA1 allele frequencies: ESRD patients vs. controls

The common alleles (percentage frequency >0.1%) identified at HLA-A, -B, -C, -DRB1, -DQB1 and DQA1 loci in ESRD patients or control population are listed in **Table 1**. We compared the percentage frequencies of these alleles between the end-stage renal disease (ESRD) patients and control group.

**Table 1:** Percentage frequencies for HLA alleles in control and ESRD Patients.

|  |  |  |  |  |  |  |
| --- | --- | --- | --- | --- | --- | --- |
| Locus | Allele | Controls (% frequency) | ESRD (% frequency) | OR | 95% CI (OR) | P- Value |
| A | A*02 | 15.92 | 14.49 | 0.708 | 0.5548-0.904 |  |
| A | A*11 | 14.88 | 13.98 | 0.930 | 0.736-1.1749 |  |
| A | A*01 | 13.76 | 12.88 | 0.918 | 0.7201-1.1694 |  |
| A | A*24 | 11.46 | 10.97 | 0.952 | 0.7336-1.2347 |  |
| A | A*26 | 9.97 | 11.27 | 1.135 | 0.8699-1.4812 |  |
| A | A*68 | 6.40 | 8.08 | 1.280 | 0.9333-1.7564 |  |
| A | A*03 | 6.18 | 7.14 | 1.169 | 0.8418-1.6224 |  |
| A | A*33 | 6.47 | 5.84 | 0.895 | 0.6354-1.2614 |  |
| A | A*31 | 5.28 | 4.63 | 0.870 | 0.5947-1.2727 |  |
| A | A*32 | 4.24 | 4.43 | 1.046 | 0.6995-1.5635 |  |
| A | A*30 | 1.71 | 2.72 | 1.604 | 0.9139-2.814 |  |
| A | A*29 | 1.71 | 1.71 | 0.999 | 0.531-1.8808 |  |
| A | A*23 | 0.89 | 0.91 | 1.014 | 0.4257-2.4165 |  |
| A | A*74 | 0.60 | 0.81 | 1.355 | 0.5068-3.6227 |  |
| A | A*66 | 0.15 | 0.10 | 0.676 | 0.0612-7.4629 |  |
| A | A*36 | 0.30 | 0.00 |  |  |  |
| A | A*66 | 0.15 | 0.10 |  |  |  |
| A | A*34 | 0.00 | 0.10 |  |  |  |
| B | B*08 | 13.84 | 14.69 | 1.072 | 0.8482-1.3547 |  |
| B | B*40 | 14.21 | 11.07 | 0.751 | 0.5848-0.9649 | p ≤ 0.05 |
| B | B*35 | 11.16 | 11.57 | 1.041 | 0.8045-1.348 |  |
| B | B*51 | 10.49 | 9.15 | 0.860 | 0.6516-1.1346 |  |
| B | B*52 | 6.85 | 6.94 | 1.015 | 0.7346-1.4029 |  |
| B | B*15 | 6.32 | 5.33 | 0.834 | 0.586-1.1877 |  |
| B | B*44 | 4.39 | 4.93 | 1.129 | 0.7661-1.6647 |  |
| B | B*07 | 4.39 | 4.53 | 1.129 | 0.7661-1.6647 |  |
| B | B*57 | 4.76 | 3.92 | 0.817 | 0.5437-1.2269 |  |
| B | B*50 | 3.20 | 4.93 | 1.569 | 1.0328-2.383 | p ≤ 0.05 |
| B | B*13 | 3.35 | 3.42 | 1.022 | 0.6499-1.6084 |  |
| B | B*58 | 3.20 | 2.92 | 0.210 | 0.1291-0.3428 |  |
| B | B*37 | 2.38 | 3.12 | 1.320 | 0.7998-2.178 |  |
| B | B*55 | 2.16 | 3.32 | 1.557 | 0.939-2.5821 |  |
| B | B*27 | 2.23 | 2.72 | 1.223 | 0.7223-2.0705 |  |
| B | B*18 | 1.56 | 1.41 | 0.900 | 0.4554-1.7788 |  |
| B | B*38 | 1.19 | 1.21 | 1.014 | 0.4777-2.1537 |  |
| B | B*41 | 0.97 | 1.21 | 1.251 | 0.5684-2.7539 |  |
| B | B*39 | 0.89 | 1.11 | 1.242 | 0.5458-2.8267 |  |
| B | B*45 | 0.67 | 0.91 | 1.355 | 0.536-3.4269 |  |
| B | B*56 | 0.52 | 0.40 | 0.772 | 0.22530-2.6435 |  |
| B | B*53 | 0.22 | 0.60 | 2.715 | 0.6772-10.8808 |  |
| B | B*49 | 0.37 | 0.30 | 0.811 | 0.1933-3.4003 |  |
| B | B*48 | 0.22 | 0.10 | 0.450 | 0.0468-4.3342 |  |
| B | B*14 | 0.15 | 0.00 |  |  |  |
| B | B*42 | 0.15 | 0.00 |  |  |  |
| B | B*68 | 0.00 | 0.10 |  |  |  |
| C | C*1 | 26.53 | 30.05 | 1.135 | 0.5806-2.2174 |  |
| C | C*2 | 15.82 | 13.79 | 1.924 | 0.4283-8.6434 |  |
| C | C*3 | 14.80 | 13.55 | 1.382 | 0.8379-2.2786 |  |
| C | C*4 | 11.90 | 11.58 | 1.049 | 0.6837-1.6107 |  |
| C | C*5 | 9.35 | 9.85 | 1.151 | 0.3071-4.3108 |  |
| C | C*6 | 5.78 | 7.88 | 0.894 | 0.6209-1.2862 |  |
| C | C*7 | 3.40 | 3.94 | 1.176 | 0.8885-1.556 |  |
| C | C*8 | 3.91 | 2.71 | 0.551 | 0.2279-1.3312 |  |
| C | C*12 | 3.06 | 2.71 | 0.843 | 0.5892-1.2071 |  |
| C | C*14 | 3.06 | 1.72 | 0.874 | 0.4085-1.8714 |  |
| C | C*15 | 0.85 | 0.99 | 0.960 | 0.6476-1.4222 |  |
| C | C*16 | 0.51 | 0.99 | 0.678 | 0.3269-1.4073 |  |
| C | C*17 | 0.85 | 0.25 | 0.286 | 0.0332-2.453 |  |
| C | C*18 | 0.17 | 0.00 | 0.000 | 0.00 |  |
| DRB1 | DRB1*3 | 20.86 | 21.23 | 1.024 | 0.8375-1.2521 |  |
| DRB1 | DRB1*15 | 19.30 | 17.51 | 0.889 | 0.7188-1.0993 |  |
| DRB1 | DRB1*7 | 12.22 | 13.18 | 1.092 | 0.8541-1.3966 |  |
| DRB1 | DRB1*11 | 10.58 | 12.17 | 1.173 | 0.9067-1.5181 |  |
| DRB1 | DRB1*13 | 9.99 | 7.55 | 0.737 | 0.5485-0.9902 | p ≤ 0.05 |
| DRB1 | DRB1*4 | 6.86 | 8.05 | 1.191 | 0.8723-1.6265 |  |
| DRB1 | DRB1*14 | 7.23 | 6.74 | 0.929 | 0.6729-1.2831 |  |
| DRB1 | DRB1*10 | 5.07 | 6.24 | 1.248 | 0.8758-1.7792 |  |
| DRB1 | DRB1*12 | 3.13 | 1.71 | 0.539 | 0.3052-0.9533 | p ≤ 0.05 |
| DRB1 | DRB1*1 | 2.24 | 2.62 | 1.176 | 0.6913-2.0021 |  |
| DRB1 | DRB1*8 | 1.12 | 1.21 | 1.083 | 0.5045-2.3234 |  |
| DRB1 | DRB1*9 | 0.60 | 1.11 | 1.859 | 0.745-4.6389 |  |
| DRB1 | DRB1*16 | 0.52 | 0.40 | 0.772 | 0.2253-2.6435 |  |
| DRB1 | DRB1*5 | 0.22 | 0.00 |  |  |  |
| DRB1 | DRB1*2 | 0.00 | 0.20 |  |  |  |
| DRB1 | DRB1*6 | 0.07 | 0.10 |  |  |  |
| DQB1 | DQB1*2 | 28.33 | 30.59 | 1.106 | 0.9237-1.3241 |  |
| DQB1 | DQB1*3 | 26.23 | 26.93 | 1.028 | 0.8539-1.2386 |  |
| DQB1 | DQB1*6 | 26.83 | 22.87 | 0.803 | 0.6629-0.9722 | p ≤ 0.05 |
| DQB1 | DQB1*5 | 17.19 | 18.29 | 1.071 | 0.8639-1.3279 |  |
| DQB1 | DQB1*4 | 1.12 | 1.22 | 1.083 | 0.5045-2.3234 |  |
| DQB1 | DQB1*1 | 0.3 | 0.1 | 0.337 | 0.0376-3.0231 |  |
| DQA1 | DQA1*1 | 44.38 | 41.87 | 0.899 | 0.7618-1.0615 |  |
| DQA1 | DQA1*5 | 33.51 | 34.45 | 1.039 | 0.8732-1.2353 |  |
| DQA1 | DQA1*2 | 10.79 | 11.89 | 1.112 | 0.858-1.4404 |  |
| DQA1 | DQA1*3 | 7.35 | 9.76 | 1.359 | 1.0127-1.8242 | p ≤ 0.05 |
| DQA1 | DQA1*6 | 2.85 | 1.12 | 0.385 | 0.1956-0.7562 | p ≤ 0.05 |
| DQA1 | DQA1*4 | 1.12 | 0.91 | 0.810 | 0.3528-1.8575 |  |

For HLA-A allele types no significant differences were observed between the ESRD patients and the control group (**Table 1**). For HLA-B allele types, only HLA-B*40 and HLA-B*50 display significant differences in their percentage frequencies between the two groups (**Table 1**). HLA-B*40 showed a significant negative association with ESRD (OR = 0.751, P ≤ 0.05) and HLA-B*50 showed a significant positive association with ESRD (OR = 1.569, P≤ 0.05). For HLA-C allele types, again no significant differences were observed between the ESRD patients and the control group (**Table 1**).

In analysis of the association of HLA-DRB1, HLA-DQB1, and HLA-DQA1 with ESRD, HLA-DRB1*13, HLA-DRB1*12, HLA-DQB1*6, HLA-DQA1*3 and HLA-DQA1*6 display significant differences in their percentage frequencies between the two groups (**Table 1**). Among these significant negative association were found for HLA-DRB1*13 (OR = 0.737, P≤ 0.05), HLA-DRB1*12 (OR = 0.539, P≤ 0.05), HLA-DQB1*6 (OR = 0.803, P≤ 0.05) and HLA-DQA1*6 (OR = 0.385, P≤ 0.05). On the other hand, a significant positive association was found only for HLA-DQA1*3 (OR = 1.359, P≤ 0.05). At each locus, few alleles were only present in either the control or ESRD population. But their allele frequencies were too low to draw any conclusion.

### Effect of heterozygosity or homozygosity of the ESRD-associated HLA alleles on prevalence and age of onset of the disease

Previous works have hypothesized that HLA polymorphism at different loci could influence the susceptibility to ESRD. Here, we aimed to determine whether heterozygosity or homozygosity of the disease-associated HLA alleles at different loci could modify the prevalence of disease. To achieve that we examined the association of different allele combinations at HLA-A, -B, -C, -DRB1, -DQB1 and DQA1 loci with ESRD. **Table 2** shows the allele combinations that displayed a significant difference in their percentage frequencies between the ESRD patients and the control group. On HLA-A locus, we identified only one allele combination A*3/A*26 that showed significant positive association (OR= 4.488, P≤ 0.05) with ESRD. Both of these alleles had separately shown a positive association with ESRD (**Table 1**); however, the differences in their percentage frequencies between control and ESRD group did not reach statistical significance. In our data, this allele showed the highest strength of association with ESRD.

**Table 2:** Percentage frequencies for HLA allele combinations in control and ESRD Patients.

|  |  |  |  |  |  |  |
| --- | --- | --- | --- | --- | --- | --- |
| Locus | Allele Combination | Controls (% frequency) | ESRD (% frequency) | OR | CI | P value |
| A | A*3_26 | 0.30 | 1.37 | 4.4881 | 0.9283-21.6984 | 0.0488 |
| B | B*40_51 | 3.55 | 1.56 | 0.4324 | 0.1918-0.9749 | 0.0434 |
| B | B*15_40 | 2.93 | 0.98 | 0.3273 | 0.1213-0.8826 | 0.022 |
| B | B*35_39 | 0.00 | 0.78 | 2.2787 | 2.1347-2.4323 | 0.0375 |
| B | B*44_44 | 0.00 | 0.78 | 2.2787 | 2.1347-2.4323 | 0.0375 |
| DQB1 | DQB1*6_6 | 9.27 | 6.07 | 0.6366 | 0.4058-0.9985 | 0.0484 |
| DRB1 | DRB1*4_13 | 2.31 | 0.58 | 0.2489 | 0.0717-0.8646 | 0.0284 |

On HLA-B locus, allele combination B*40/B*51 showed significant negative association (OR = 0.4324, P≤ 0.05 with ESRD. In the initial analyses, B*40 showed a significant negative association with ESRD (**Table 1**). B*51 allele also showed a negative association with ESRD (**Table 1**); however, the differences in its percentage frequency between control and ESRD population did not reach statistical significance. The combination B*40/B*15 showed a significant negative association (OR = 0.3273, P≤ 0.05) with ESRD. When analysed separately, the B*15 allele also showed a negative association with ESRD (**Table 1**); however, its percentage frequency was not significantly different between control and ESRD population. Homozygous combination B*44/B*44 and heterozygous combination B*35/B*39, were only present in ESRD patients but the percentage frequencies for these combinations were too low to conclusively claim their significance.

At HLA-DQB1 locus, a homozygous combination of DQB1*6/ DQB1*6 showed a significant negative association (OR = 0.6366, P<0.05) with ESRD. At HLA-DRB1 locus, a combination of DRB1*4/DRB1*13 showed a significant negative association (OR = 0.2489, P<0.05) with ESRD.

Next, we sought to determine whether different HLA allele combinations affect the age of onset for ESRD. **Supplementary Figure 1** compares the age of onset for all the identified allele combinations (percentage frequency > 0.1 %) at HLA-A, -B, -C, -DRB1, -DQB1 and DQA1 loci. We observed certain differences in age of onset among individuals carrying different allele combinations. However, these differences fail to reach statistical significance.

## Discussion

Human leukocyte antigen (HLA) has been associated with a variety of renal disorders [1] and has emerged as a significant risk factor in most immune-mediated renal diseases. HLA associations have also been described for renal diseases that are less traditionally seen as autoimmune or immune-mediated [1]. Several works have studied possible associations between HLA types and ESRD [3-5]. Most of these reports have studied waitlisted candidates for renal transplantation in homogeneous ethnic populations. Although underpowered, most of these studies have not defined significant associations and to date, no consistent HLA associations with ESRD itself have been identified. The major aim of the presented work was to revisit the association between HLA alleles and ESRD in the Pakistani population and compare our data with previous major works. As shown above, our study also identified multiple alleles and allele combinations that showed significant associations with ESRD. We also performed an extensive literature survey and collected information on significant HLA and ESRD associations from previous studies. We aimed to identify any consistent HLA associations with ESRD that can be similar across different ethnic groups. **Supplementary Table S2** shows the HLA allele types that have shown significant associations (negative or positive) with ESRD in our work or previous studies. For HLA-A locus, the HLA-A*11 allele showed a significant positive association with ESRD in three separate studies –that included study-participants from different populations [12-14], while, HLA-A*28 showed a significant negative association with ESRD in two of the previous works [15, 16]. Other HLA-A alleles also showed significant associations with ESRD. However, the association was noted in only one study or there were contradictions in the trends of association (negative or positive) in separate works. For HLA-B locus, multiple alleles showed significant associations with ESRD. HLA-B*15 was found to be positively associated with ESRD by four separate works [12, 14, 17, 18]. HLA-B*55[14] [19] and HLA-B*53 [13, 20] were observed to be positively associated with ESRD in two studies. On the other hand, HLA-B*52 was negatively associated with ESRD in two studies [13, 21]. HLA-B*40 and HLA-B*50, that respectively showed a significant negative and positive association with ESRD in our study, showed contradictory but significant trends of associations by other works. HLA-B*18, HLA-B*39, and HLA-B*8 also showed contradictions in the trend of association (negative or positive) in separate works. Multiple alleles at HLA-C locus also showed significant associations with ESRD but we do not find any consistent patterns among different studies.

Several HLA-DRB1 alleles also displayed consistent patterns of association with ESRD in different populations, for instance, HLA-DRB1*3, *4 and *11 all showed a significant positive association with ESRD in several separate works (**Supplementary Table S2**). HLA-DQB1*6 allele was found to be negatively associated with ESRD in our study as well as three other works. Multiple other alleles also showed significant associations with ESRD. However, these observations had certain limitations, for instance, the association was noted in only one study or there were contradictions in the trend of association (negative or positive).

Hence, these comparisons between our data and previous works show that at least for certain alleles, the association of HLA to ESRD can be similar across different ethnic groups. It has been previously hypothesized that the association of HLA types with ESRD might be confounded by the presence of HLA associations with other diseases. For instance, a large, phenome-wide association study (PheWAS), showed that the HLA-DQB1*03:02 allele is associated with kidney transplantation (OR 1.4) –which was potentially related to the increased risk of and diabetic kidney disease (OR 7.1) [22]. Few studies suggest that HLA types can affect the severity of kidney disease or risk of progression independent of disease etiology [1]. The comparison of HLA types among ESRD patients and a control group without detailed clinical phenotyping might miss some of the important but subtle potential effects of HLA types, including the influence of HLA on the age of disease onset, disease severity, disease phenotype and rates of disease triggering events. Further studies are required to study the effect of HLA types on the aforementioned factors.

## Abbreviations

ESRD: End-Stage Renal Disease
HLA: Human Leukocyte Antigen

## Acknowledgments

We would like to acknowledge Mr. Banaras Masih for his technical assistance in HLA genotyping of the samples.

---

**Supplementary Table S1.**
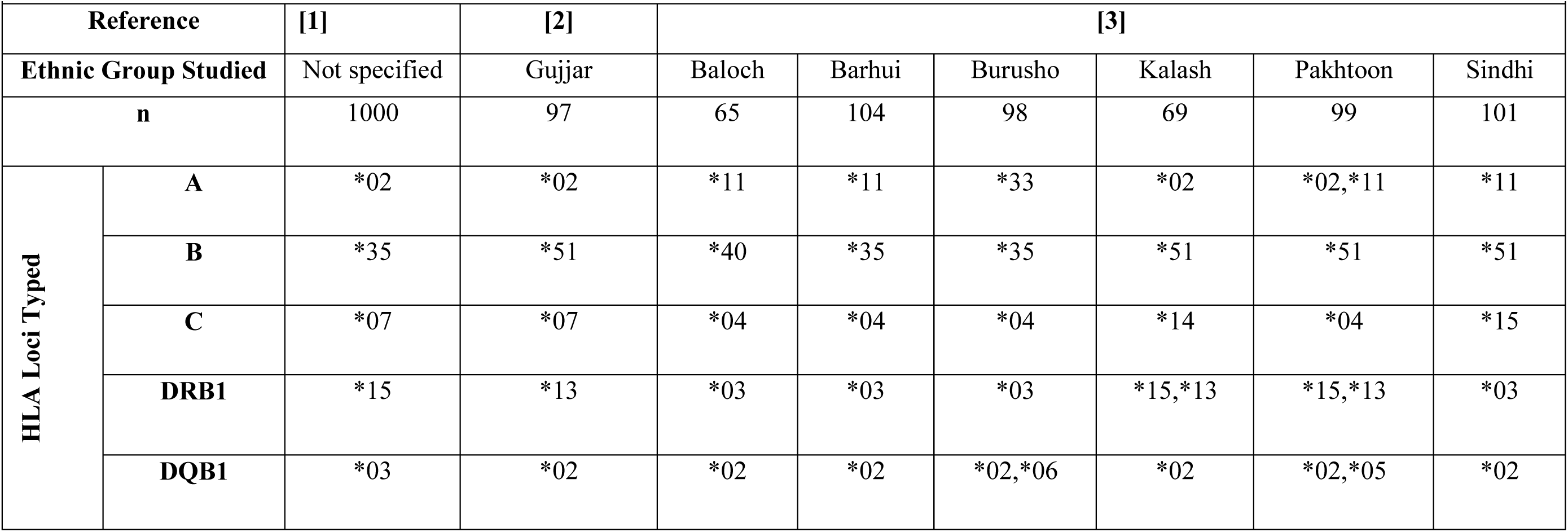
Most frequent HLA alleles in Pakistani population: data from previous works.

|  |  |  |  |  |  |  |  |  |  |
| --- | --- | --- | --- | --- | --- | --- | --- | --- | --- |
| Reference |  | [1] | [2] | [3] |  |  |  |  |  |
| Ethnic Group Studied |  | Not specified | Gujjar | Baloch | Barhui | Burusho | Kalash | Pakhtoon | Sindhi |
| n |  | 1000 | 97 | 65 | 104 | 98 | 69 | 99 | 101 |
| HLA Loci Typed | A | *02 | *02 | *11 | *11 | *33 | *02 | *02,*11 | *11 |
|  | B | *35 | *51 | *40 | *35 | *35 | *51 | *51 | *51 |
|  | C | *07 | *07 | *04 | *04 | *04 | *14 | *04 | *15 |
|  | DRB1 | *15 | *13 | *03 | *03 | *03 | *15,*13 | *15,*13 | *03 |
|  | DQB1 | *03 | *02 | *02 | *02 | *02,*06 | *02 | *02,*05 | *02 |

**Supplementary Table S2:**
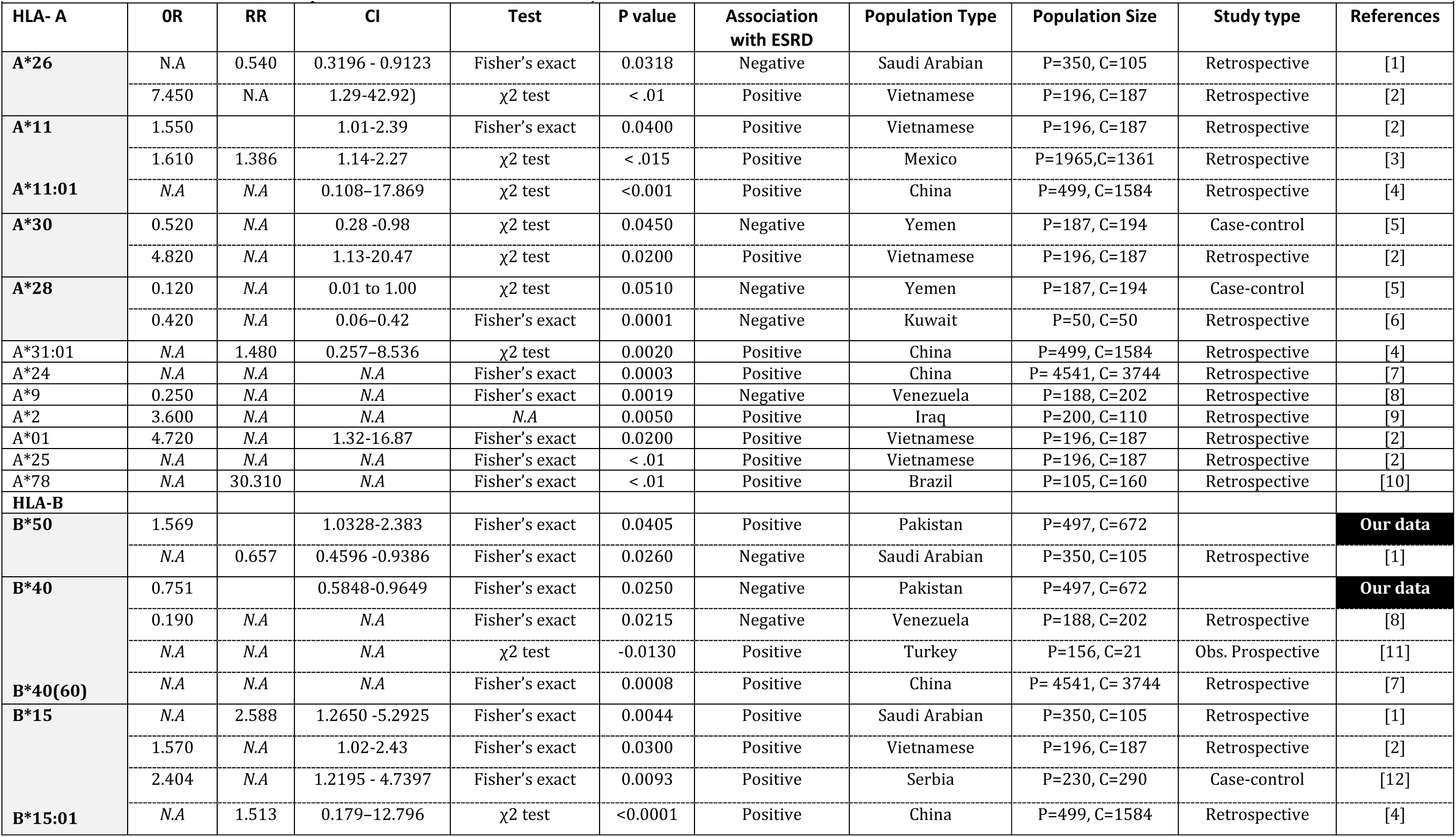
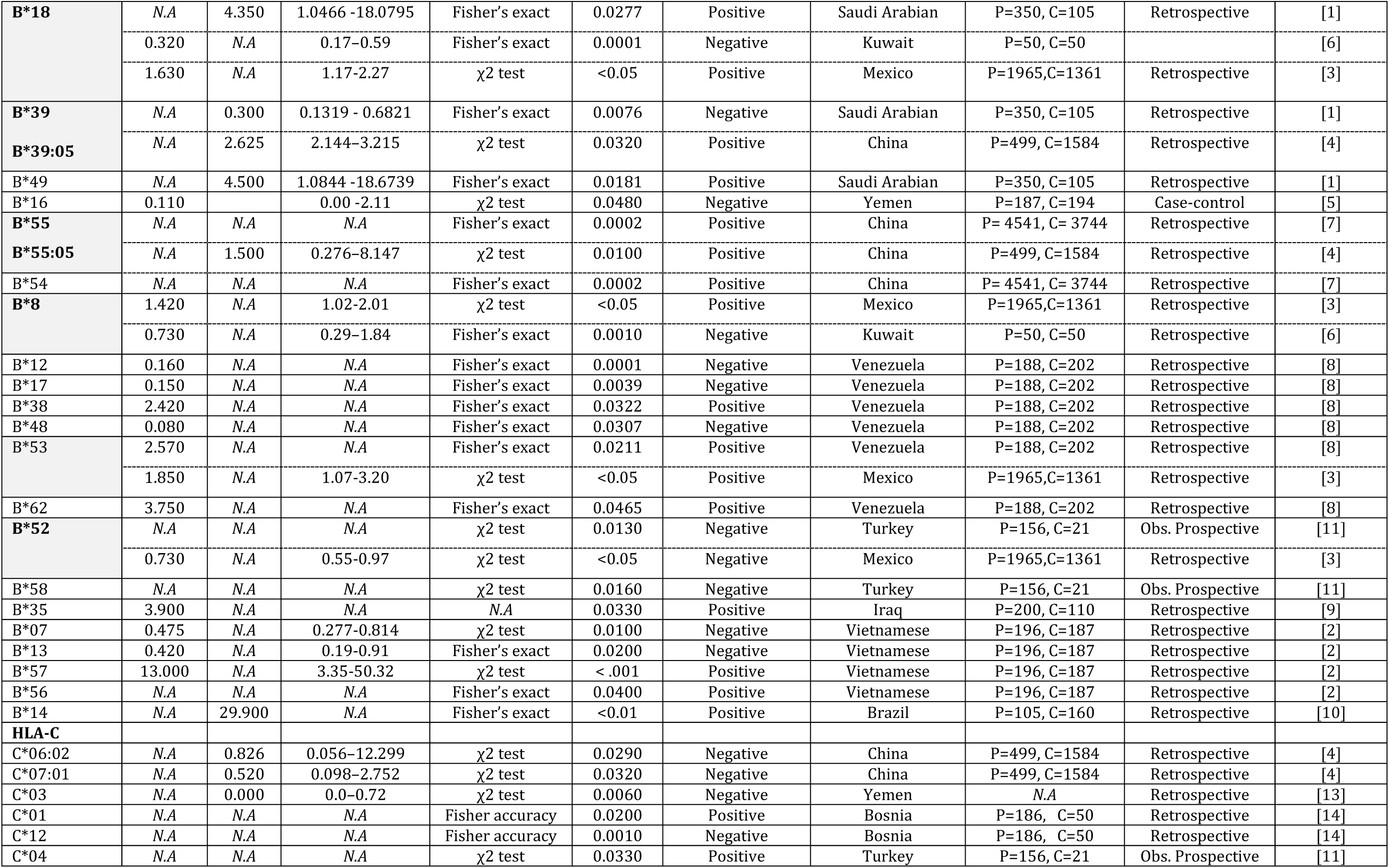
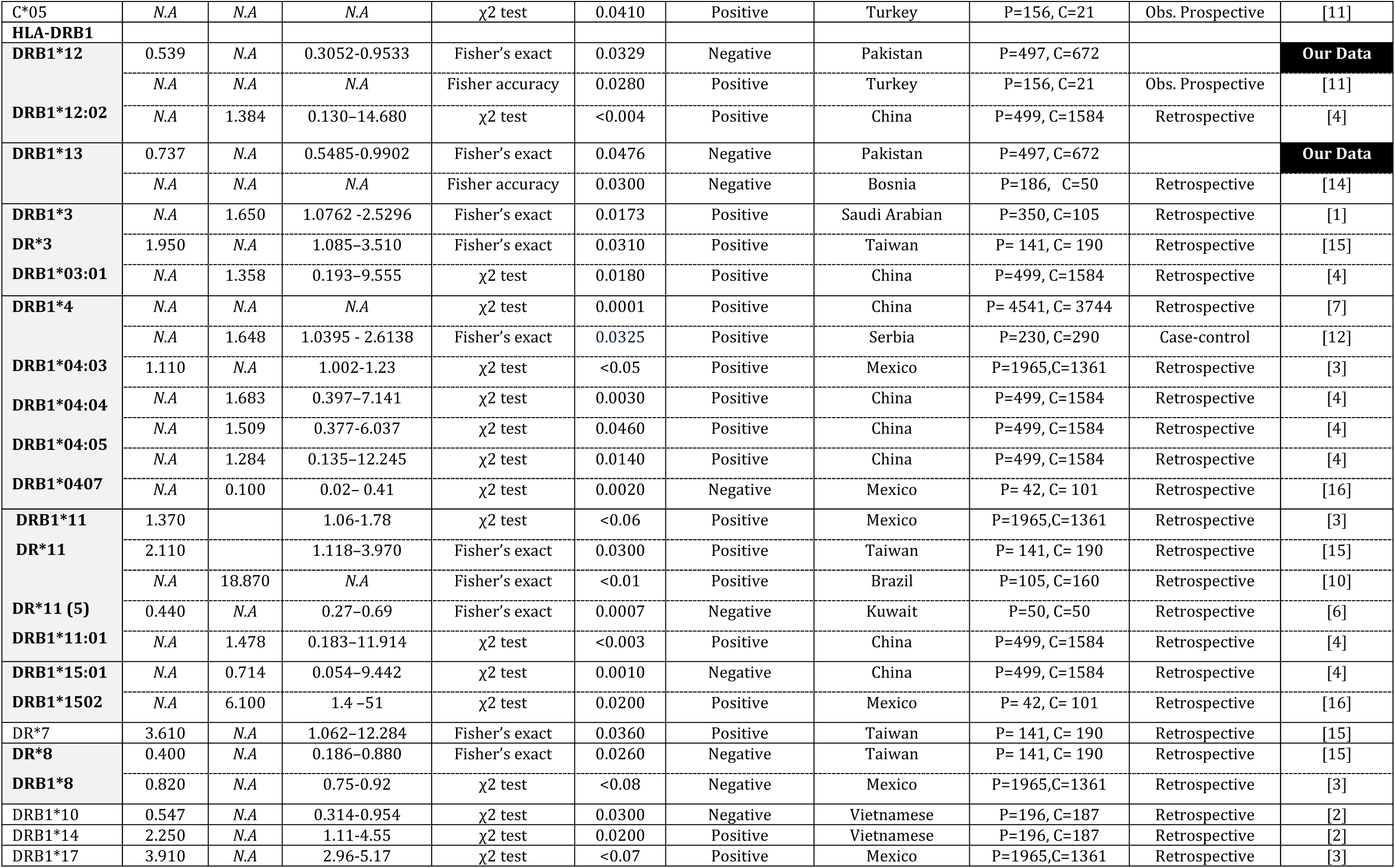
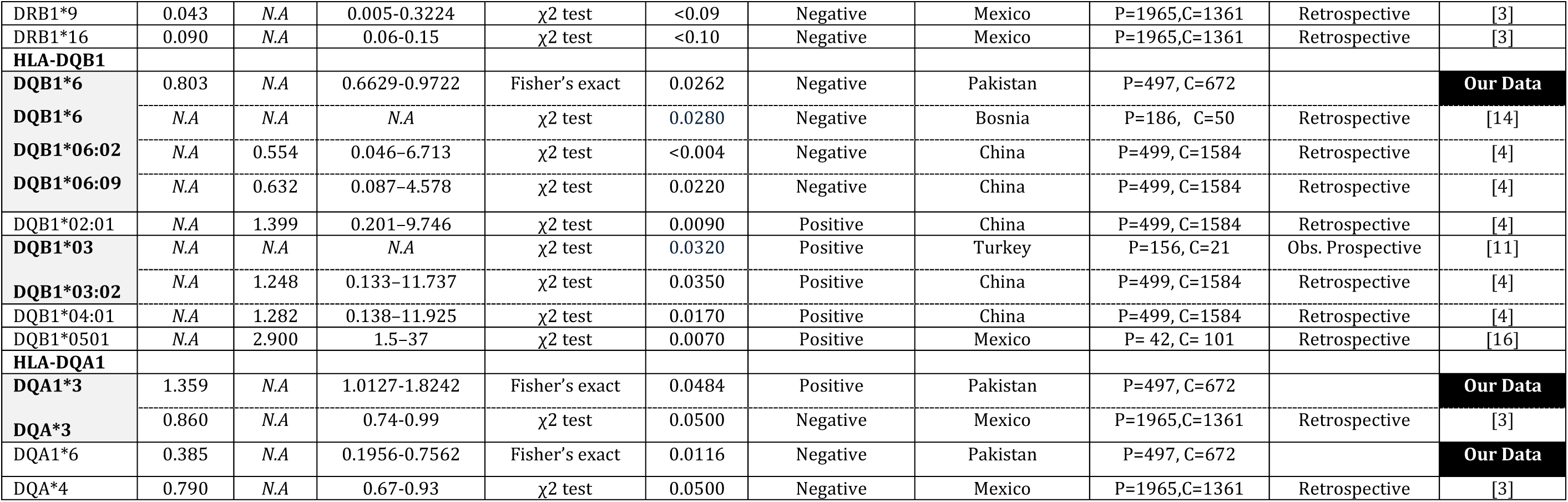
HLA alleles and ESRD. To identify any consistent association of HLA alleles with ESRD, we performed an extensive literature survey and collected information on significant associations. The allele types that displayed significant associations in more than one study are highlighted (**Bold Letters**/Grey Cells). **Abbreviations:** P, number of ESRD patients; C, number of control subjects; N.A, Not Available.

**Supplemtary Figure 1:**
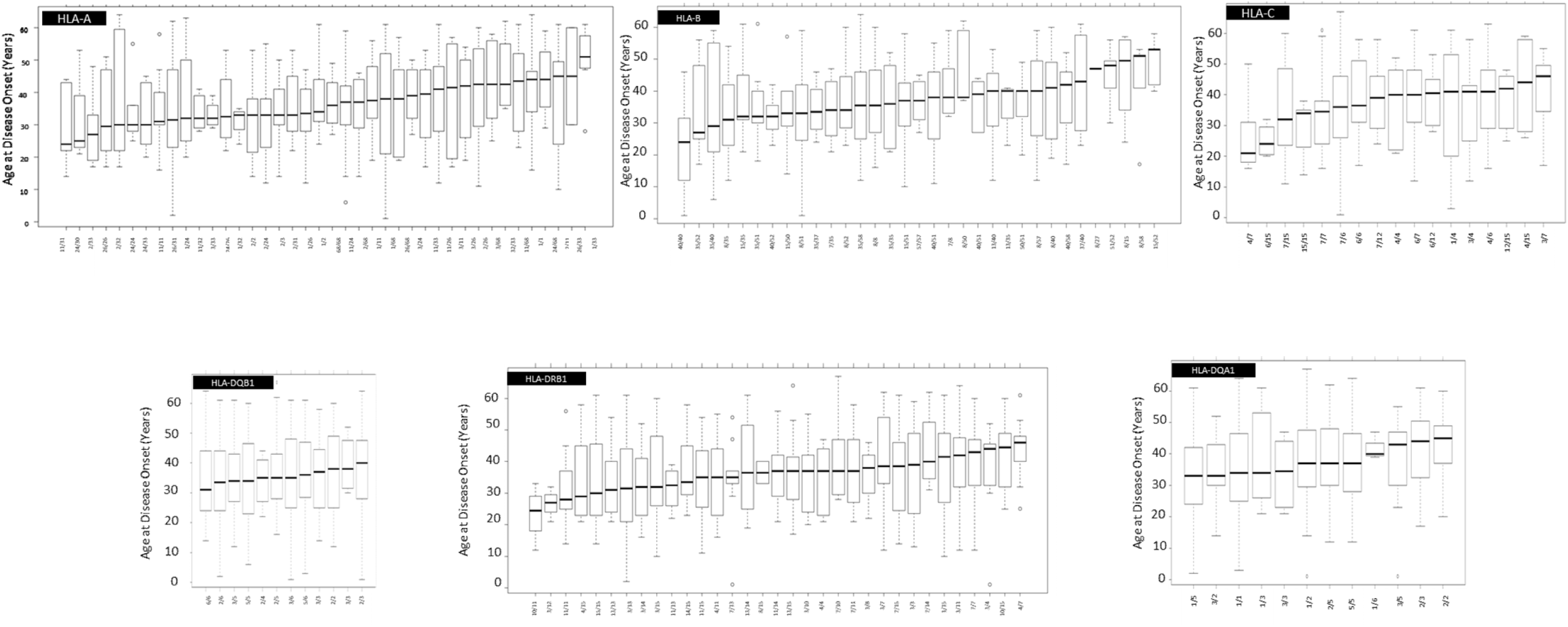
Effect of allele combinations at HLA-A, -B, -C, -DRB1, -DQB1 and DQA1 loci on the age of onset for ESRD. The ANOVA test, complemented by a pairwise Tukey’s HSD test, is used to examine significant differences among the groups, with a *P* value threshold of 0.05.

